# Extracellular vesicles from preeclampsia disrupt the blood-brain barrier via reduced claudin-5: potential role of vascular endothelial growth factor

**DOI:** 10.1101/2024.04.10.588960

**Authors:** Hermes Sandoval, Belén Ibañez, Moisés Contreras, Felipe Troncoso, Fidel O Castro, Diego Caamaño, Lidice Mendez, Estefanny Escudero-Guevara, Hiten D. Mistry, Lesia O. Kurlak, Manu Vatish, Jesenia Acurio, Carlos Escudero

## Abstract

**Background:** Physiopathology of life-treating cerebrovascular complications in preeclampsia are yet unknown. We investigated whether disruption of the blood-brain barrier (BBB), generated using circulating small extracellular vesicles (sEVs) from women with preeclampsia or placentae cultured under hypoxic conditions, impairs the expression of tight junction proteins, such as claudin 5 (CLDN5), mediated by VEGF and activation of VEGF receptor 2 (KDR).

**Methods:** sEVs were isolated from plasma (normal pregnancy, sEVs-NP, n=9); preeclampsia, sEVs-PE, n=9) or placental explants from normotensive pregnancies, cultured in normoxia (sEVs-Nor, n=10) or hypoxia (sEVs-Hyp, n=10). The integrity of the BBB was evaluated using *in vitro* (human and mice brain endothelial cell lines) and *in vivo* (non-pregnant C57BL/6 mice (4 to 5 months old, (n=10) were injected with sEVs-Hyp), models.

**Results:** sEVs-PE and sEVs-Hyp reduced CLND5 levels (p<0.05) in the endothelial cell membrane without affecting other tight junction proteins. These results were negated with sEVs-PE sonication. sEVs-Hyp injected into non-pregnant mice generated neurological deficits and BBB disruption, specifically in the posterior area of the brain, associated with reduction in CLND5 levels in the brain cortex. Furthermore, sEVs-PE and sEVs-sHyp had higher VEGF levels than sEVs-NP and sEVs-Nor, respectively. Human brain endothelial cells exposed to sEVs-PE or sEVs-sHyp exhibited a reduction in the activation of KDR.

**Conclusion:** sEVs from hypoxic placentae and plasma from women with preeclampsia disrupt the BBB, via reduction of CLDN5, a phenomenon that may involve VEGF contained within these vesicles. These findings will improve the elucidation of cerebrovascular alterations in women with preeclampsia.

## Introduction

Preeclampsia (PE) is one of the leading causes of maternal mortality in low and middle-income countries (LMIC)^1^. Most maternal deaths associated with PE are related to cerebrovascular complications, which include epilepsy, stroke, or posterior reversible encephalopathy syndrome (PRES), among others^2-4^. The physiopathology of the cerebrovascular complication in PE remains unclear. However, we and other groups have reported disruption of the blood-brain barrier (BBB), generated by hitherto unidentified circulating factors in the maternal circulation^5-8^. Interestingly, we recently reported that small extracellular vesicles (sEVs), derived from the placentae cultured in hypoxia or from plasma of women with PE, disrupt the BBB ^7^. However, it is unclear which factor(s) within the sEVs might be responsible for this disruption in PE: candidates such as proinflammatory cytokines^8^ or vascular modulators (such as the vascular endothelial growth factor; VEGF)^5,9^ have been independently suggested to play a role.

As a critical BBB component, the brain endothelial cells exhibit a highly specialized phenotype with four key functional characteristics: high expression of tight junction proteins; several specific influx and efflux transporters; low vesicular transport and a distinct glycocalyx with low expression of adhesion molecules^10-12^. Among them, tight junction proteins (TJs) create an apical intercellular junctional complex that works as a barrier to paracellular diffusion of blood-borne polar substances, maintaining cell polarization, but also acting as an intracellular signaling platform^13,14^. Thus, brain endothelial cells have a high expression of claudin-5 (CLDN5) (mRNA levels of CLDN5 are approximately 600 times higher compared to other CLDNs)^15,16^. Moreover, deletion of CLDN5 is life-treating since all CLDN5 deficient mice die within ten hours after birth^17^.

Little is known regarding endothelial cell expression of CLDN5 in PE. Liévano et al.,^18^ reported reduced protein levels of CLDN5 in placental endothelium from women with PE compared with normotensive pregnancies. In the brain, ShamsEldeen et al.,^19^ showed that PE-like syndrome in rats (i.e., generated by oral administration of nitric oxide inhibitor) is associated with reduced brain protein levels of CLDN5 and high-water content of the brain.

Interestingly, recent evidence from preclinical models of acute ischemic stroke has found that sEVs derived from M2 microglia^20,21^ or brain endothelial cells^22^ can reduce the damage of the BBB, increasing the expression of CLDN5 in brain tissue. However, there are not information whether circulating or placenta-derived sEVs may regulate brain levels of CLND5, neither regarding its potential implication in the disruption of the BBB observed in PE.

Therefore, we hypothesize that disruption of the BBB generated by sEVs isolated from placentas cultured in hypoxia (sEVs-Hyp) or from plasma of women with PE (sEVs-PE) involves the reduction in the CLDN5 protein levels in brain endothelial cells. We also hypothesize that this reduction in CLDN5 is associated with the VEGF cargo in the sEVs-Hyp or sEVs-PE.

## Methods

Human circulating sEVs were isolated from heparinized plasma of pregnant women with normal pregnancy (NP; n=9) and preeclampsia (PE; n=9), defined as we previously reported^7^. Table S1 includes the clinical characteristics of the patient sub-group in this manuscript.

### Animals

Animal studies were approved by the ethics and biosafety committee of the Universidad del Bio Bio under ethical principles of humanized animal management and “Three R” (3R’) principles (Fondecyt grant 1240295). Non-pregnant female C57BL/6 mice (4-6 months of age) were included. These animals were kept under standard conditions, as reported previously^23^, which included relative humidity (40-60%), room temperature (18-24°C), and photoperiod (12 h light / 12 h dark). Regular food, such as Rodent Diet 5001 (Labdiet®CA) and tap water, were *ad libitum*.

Three experimental groups of mice were generated. 1) control group (CTL, intravenous injection of 0.9% NaCl solution; n=10); 2) mice treated with sEVs (200 μg total sEVs protein), isolated from placentae cultured in normoxic conditions (sEVs-Nor; 8% O_2_; n=10); and 3) mice treated with sEVs (200 μg total sEVs protein) isolated from placentas cultured in hypoxic conditions (sEVs-Hyp; 1% O_2_; n=10). Protocols for placental sEVs isolation, sEVs characterization^7^ and animal injection were recently published by our group^24^. Animals were randomly assigned to each group, and clinical evaluation was blinded to the experimental group.

As a comparative model, the surgical model of the PE-like syndrome in mice-sRUPP (Selective Reduced Uterine Perfusion Pressure; n=3), was performed as previously reported ^25^, to evaluate BBB disruption in this PE model and its respective sham controls (n=3; See supplementary methods).

### Analysis of Rapid Murine Coma and Behavior Scale (RMCBS)

The presence of neurological clinical syndrome in mice was assessed using the RMCBS^26^, before and after injection of the sEVs. The RMCBS evaluates ten parameters, each with a score of 2. The maximum score of 20 was considered a full-normal evaluation. Each animal was randomly analyzed within 3 minutes, without anesthesia, in two 90-second periods, with minimal stress due to the operator intervention. Additionally, all the experiments were generated in a head-to-head comparison. This evaluation was carried out at 0, 3, 6, 12, and 24 h after injection of sEVs.

### sEVs isolation

sEVs were obtained from a conditioned medium of explants of NP placentae cultured in normoxia (n=10) or hypoxia (n=10), and from plasma of women with NP (n=9) or PE (n=9) as reported previously^7^. In this manuscript, sEVs isolates from plasma were characterized by size, concentration, and protein markers (i.e., HSP70, CD81, CD63, Alix, TSG101). Placental phosphatase alkaline (PLAP) was used as a marker of the placental origin of sEVs. The sEVs content of angiogenic modulators, including VEGF, placental growth factor (PlGF), and VEGF receptor 2 (KDR), were estimated by Western blot.

### sEVs uptake

PKH67 Green Fluorescent Cell Linker Mini Kit (MINI67-1KT, Sigma-Aldrich, MO, USA), was used to fluorescently label the sEVs based on a protocol previously published^7^. Briefly, 100 μg/ml of sEVs pre-treated with the fluorescent dye was applied to hCMEC/D3 monolayers for 30 minutes at 37°C or 4°C. The cells were then fixed with paraformaldehyde (PFA 4%) and kept at 4°C prior to microscopic fluorescent analysis (Microscope Motic Scientific, USA).

In parallel experiments, sEVs (200 μg) labeled with PKH67 from the sEVs-Nor and sEVs-Hyp groups were injected into non-pregnant mice (n=2 per group), as indicated above. The CTL group was also incorporated into this analysis (n=2). 30 min after injection, animals were sacrificed for brain extraction. The brain was fixed with PFA 4% (4°C, 48 h), then cut coronally (50 μm thickness) and visualized under a confocal microscope.

### Placenta-derived sEVs *in vivo* injection

*In vivo* injection of sEVs-Nor and sEVs-Hyp preparations (200 μg of total sEVs protein) was previously reported^24^. Under anesthesia, sEVs solutions were aseptically injected with a syringe (30-gauge needle) into a mouse’s external jugular vein. The contralateral external jugular vein was sampled 6 and 24-hour post-sEVs-injection and serum produced by centrifugation (1,400 rpm x 10 minutes). Subsequently, serum samples were stored at -80°C until Western blot analysis of VEGF, PlGF, PLAP, Interleukin 6 (IL-6), Interleukin 8 (IL-8), and tumor necrosis factor-alpha (TNF-α).

### Inflammatory protein analysis in serum. Dot blot

Circulating proteins associated with angiogenesis (VEGF, PlGF, and VEGFR1), inflammation (IL-6, IL-8, and TNF-a) were identified 6 and 24 hours post injection (sEVs-Nor and sEVs-Hyp) using dot blot assay^27^. Ten μg (final volume 2 μl for all samples) of serum proteins were loaded onto a nitrocellulose membrane (Bio-rad Laboratories, CA, USA). Then, protein identification was performed using the respective antibody as indicated for Western blot (Table S2). Ponceau Red staining was used as a loading control.

### Evan’s blue extravasation analysis

Evan’s blue extravasation was evidenced in each brain section using Image J, as described previously^7,24^.

### Western blot analysis for CLDN5

CLDN5 protein expression was analyzed *in vitro* and *ex vivo* samples. For *in vitro* measurements, the human (hCMEC/D3) and mouse (BEND/3) brain endothelial cell lines were treated with 100 μg/ml of sEVs extracted from placentae or plasma, as indicated above. For *ex vivo* analysis, the brain sections were extracted for tissue homogenates and protein determination as indicated above, after mice were injected with sEVs-Nor, sEVs-Hyp (6 h).

From those samples, 50 μg of total protein extraction was used for semiquantitative analysis of CLDN5 (Santa Cruz Biotechnology, sc-374221, TX, USA). Then, CLDN5 was identified with an anti-mouse IgG secondary antibody conjugated with peroxidase (Sigma-Aldrich, MO, USA). Western blot signal was detected with a GeneGnome XRQ fluorescent detector (Syngene, CA, USA), and densitometry of CLND-5/β-actin ratio was estimated using the ImageJ 1.48 software (National Institute of Health, USA). Other tight junction proteins were also analyzed, as indicated in supplementary information (see supplementary information and Table S2).

### Cell fractionation

After total protein extraction of hCMEC/D3 cells stimulated with sEVs-Nor and sEVs-PE, as indicated above, were passed through a syringe (23 gauge) 15 times and then centrifuged at 2,000 rpm x 10 min. The supernatants were extracted and transferred to a new vial. Then, it was centrifuged again (14,000 rpm x 30 min) to obtain the cytoplasm (supernatant) and membrane (pellet) fractions. The pellet (i.e., membrane fraction) was diluted in the extraction protein buffer. With these samples, we analyzed the protein levels of CLDN5 by Western blot. Ponceau red staining was used as a loading control.

### sEVs sonication

sEVs (100 μg, diluted in 200 μL of basal medium of hCMEC/D3) of sEVs-NP and sEVs-PE were broken down by sonication (7 times of 1 min-cycles) (Cleaner Ultrasonic Set, Daiham Scientific Co, Korea). Subsequently, sonicated sEVs were used to treat hCMEC/D3 cells as previously indicated. Cell extractions were then used for CLDN5 protein amount analysis using Western blot, as indicated above.

### Statistical analysis

Quantitative variables are presented as mean ± standard deviation, whereas qualitative variables are presented in percentages considering their respective group. Non-parametric analysis was used to compare differences between groups. All studies were performed head-by-head, and data was normalized concerning the control group (i.e., injected with vehicle phosphate buffer solution, PBS). P<0.05 was considered a statistically significant difference. Data and statistical analyses were performed using the Microsoft Excel database and GraphPad Prism 6 (GraphPad Software, CA, USA).

## Results

### Characterization of sEVs

Isolates of sEVs from the plasma of women with PE (sEVs-PE) and NP (sEVs-NP) and placentae cultured in normoxia (sEVs-Nor) or hypoxia (sEVs-Hyp) are shown in Figure 1. These sEVs were characterized by protein markers, including HSP 70, CD81, Alix, TSG101, and CD63 (Figure 1A), as well as the presence of PLAP identifying the placental source of some sEVs. There were no statistical differences between the particle concentration or the size (144.0 ± 27.3 versus 153.1 ± 34.7 nm, respectively) of the vesicles between sEVs-PE and sEVs-NP (Figure 1B).

**Figure 1.**
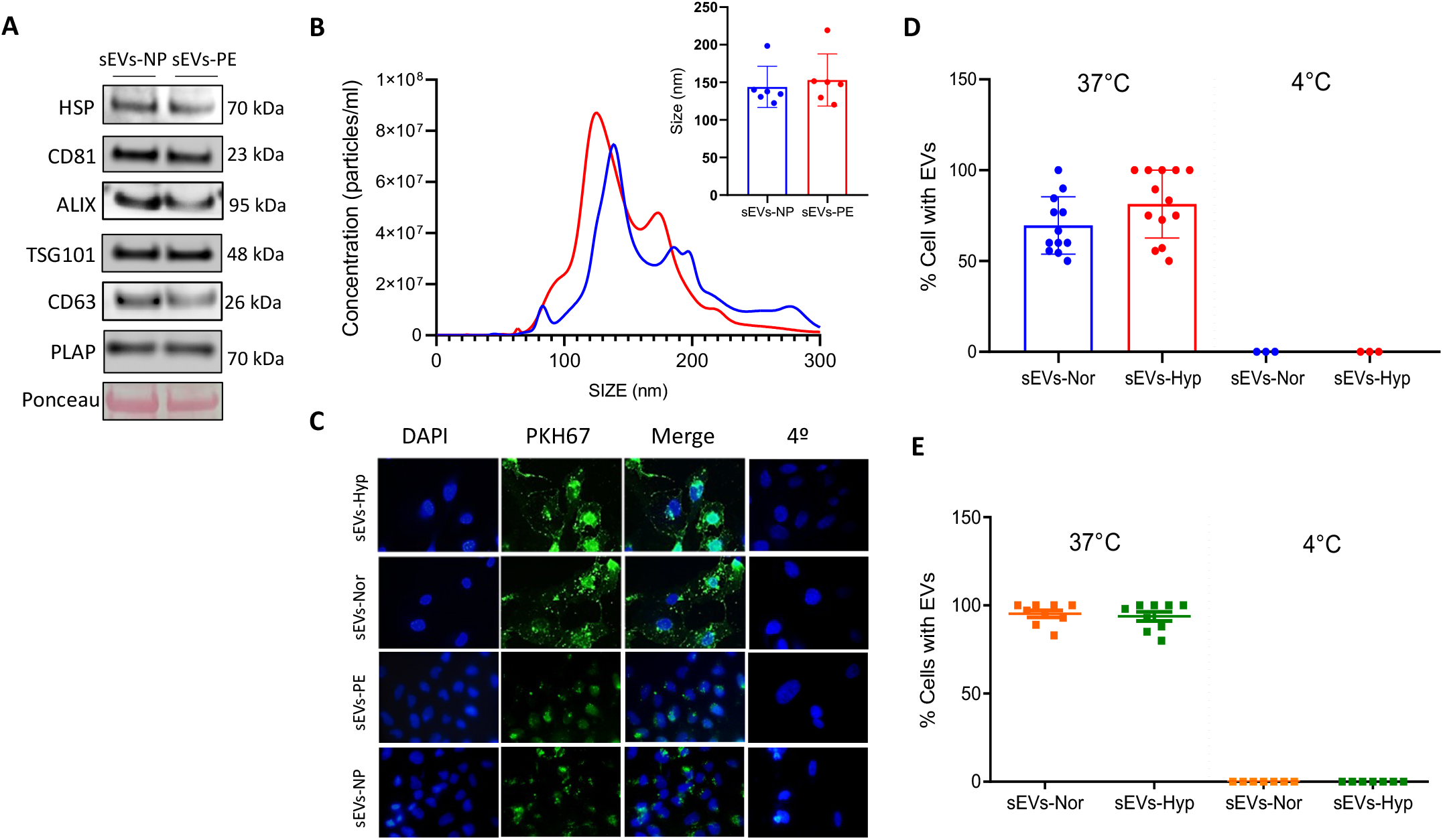
Characterization and cell internalization of sEVs. **A)** sEVs were isolated from plasma of normal pregnancy (sEVs-NP, blue), plasma from women with preeclampsia (sEVs-PE, red), placentae from normal pregnancy cultured under normoxia (8% O_2_, sEVs-Nor, orange) and placentae from normal pregnancy cultured under hypoxia (1% O_2_, sEVs-Hyp, green). Representative images of sEVs marker (HSP70, CD81, Alix, TSG101, CD63) or placental marker (PLAP). Loading control was Ponceau Red staining. **B)** Size and concentration distribution of sEVs-NP and sEVs-PE. The inserted chart that was inserted indicates the sEV size in both groups. **C)** Incorporation of sEVs loaded with PKH67 in human brain endothelial cells. Experiments were performed under at standard culture conditions (37 °C) and 4 °C. **D)** Percentage of cells that uptake sEVs-Nor and sEVs-Hyp; **E)** or sEVs-NP or sEVs-PE at 37 °C or 4 °C. Every dot represents an individual sEV extraction from an individual patient. Data are presented as mean ± SD.

Figure 1C shows representative images of the sEVs-NP, sEVs-PE, sEVs-Nor, and sEVs-Hyp uptake by human brain endothelial cells (hCMEC/D3). The percentage of cells that uptake the sEVs was estimated at 37°C or 4°C. The former estimates the active cellular incorporation of the vesicles (Figure 1D and 1E). There were no statistical differences in the sEVs uptake among the studied groups of sEVs (P>0.05).

### sEVs-PE and sEVs-Hyp reduce CLDN5 *in vitro*

We estimated the expression of the tight junction proteins in hCMEC/D3 in the absence or in the presence (0 to 6 h) of plasma or placental-derived sEVs. Cells exposed (3 h) to sEVs-PE showed a significant reduction in the protein level of CLDN5 (Figure 2A-2B; P=0.02), without changes in the expression of other tight junction proteins, including OCL-1, ZO-1 (Figure S1). Furthermore, compared with sEVs-Nor, a reduction in CLDN5 was also found after treatment with sEVs-Hyp (Figure 2C; P=0.007). To extend these *in vitro* findings to an *in vivo* setting, we analyzed CLDN5 levels in brain sections of the PE-like syndrome model, i.e., RUPP^25^. Compared with sham pregnant mice, RUPP dams exhibited disruption of the BBB, mainly in the posterior areas of the brain (Figure S2A). This effect was associated with a lack of compensatory increase in the CLDN5 in the posterior area of the brain in the RUPP dams, as was found in the sham group (Figure S2).

**Figure 2.**
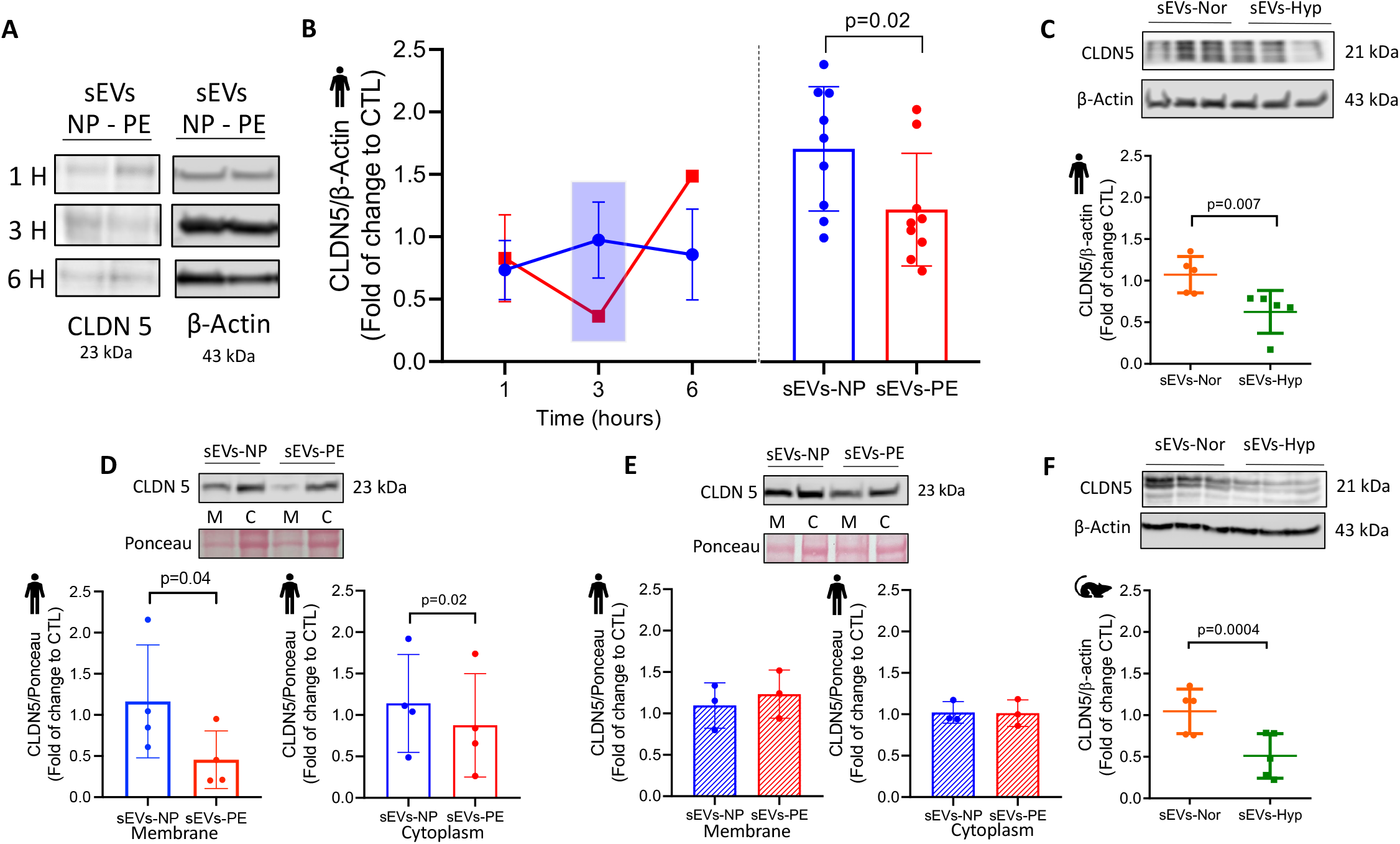
CLDN5 protein levels in brain endothelial cells treated with sEVs-PE or sEVs-Hyp. **A)** Representative blots of CLDN5 protein levels in human brain endothelial cells treated (1, 3, and 6 h) with sEVs-NP (blue bars) or sEVs-PE (red bars). β-actin was used as a loading control. **B)** Densitometry analysis of CLDN5/β-actin ratio expressed as fold of change respect to control (CTL, treated with sEVs vehicle). Remarked results at 3 hours are presented individually in the inserted chart. **C)** Representative blot and CLDN5/β-actin ratio as in B in human brain endothelial cells treated with sEVs-Nor (orange dots) or sEVs-Hyp (green dots). **D)** CLDN5 levels in membrane (M) and cytoplasmatic (C) fractions from human endothelium. Ponceau Red was used as a loading control. **E)** CLDN5 levels in M and C fractions as in D using sonicated sEVs. **F)** CLDN5 levels in mouse brain endothelial cells treated with sEVs-Nor or sEVs-Hyp. Every dot represents an individual sEVs extraction from an individual patient. Data are presented as mean ± SD. P value is included when significant differences were found.

To further explore the effect of sEVs-PE on CLDN5 cell localization, we extracted cytoplasmic and membrane fractions from hCMEC/D3 exposed (3 h) to sEVs. Again, a significant reduction in the CLDN5 levels was observed in both membrane (P=0.04) and cytoplasmatic (P=0.02) fractions in cells exposed to sEVs-PE compared with those exposed to sEVs-NP (Figure 2D). This reduction in CLDN5 in fractionated cell extractions required the integrity of sEVs-PE since hCMEC/D3 treated with sonicated sEVs-PE did not show any significant changes in the CLDN5 membrane or cytoplasmatic levels (Figure 2E).

In parallel experiments and considering our previous findings in non-pregnant mice injected with sEVs-Hyp^7^, we found that sEVs-Hyp also reduced the protein levels of CLDN5 *in vitro* in mice brain endothelial cells (Bend3; Figure 2F; P=0.0004).

### sEVs-Hyp disrupts the BBB via the reduction of CLDN5 *in vivo*

To evaluate the *in vivo* effects of sEVs-Hyp over CLDN5, we first characterized the effect of these vesicles on the neurological status of non-pregnant mice (Figure 3A). sEVs-Hyp decreased the neurological score in receiver mice, showing statistically significant differences after injection from 6 h until 24 h (end of the experiment), compared with controls treated with sEVs-Nor (Figure 3B). This reduction was more evident in the coordination and strength/tone parameters of the RMCBS (Figure 3C). The harmful neurological effect of sEVs-Hyp was evident despite no changes in the circulating levels of proinflammatory cytokines, such as IL-6, IL-8, and TNF-α 6 hours after injection (Figure 3D-3F). However, these were elevated 24 h after sEVs-Hyp injection (Figure S3). In addition, the circulating levels of PlGF (Figure S4B) or PLAP (Figure S4C), were also significantly elevated, without changes in the circulating levels of VEGF in the sEVs-Hyp injected mice after 6 h of injection (Figure S4A).

**Figure 3.**
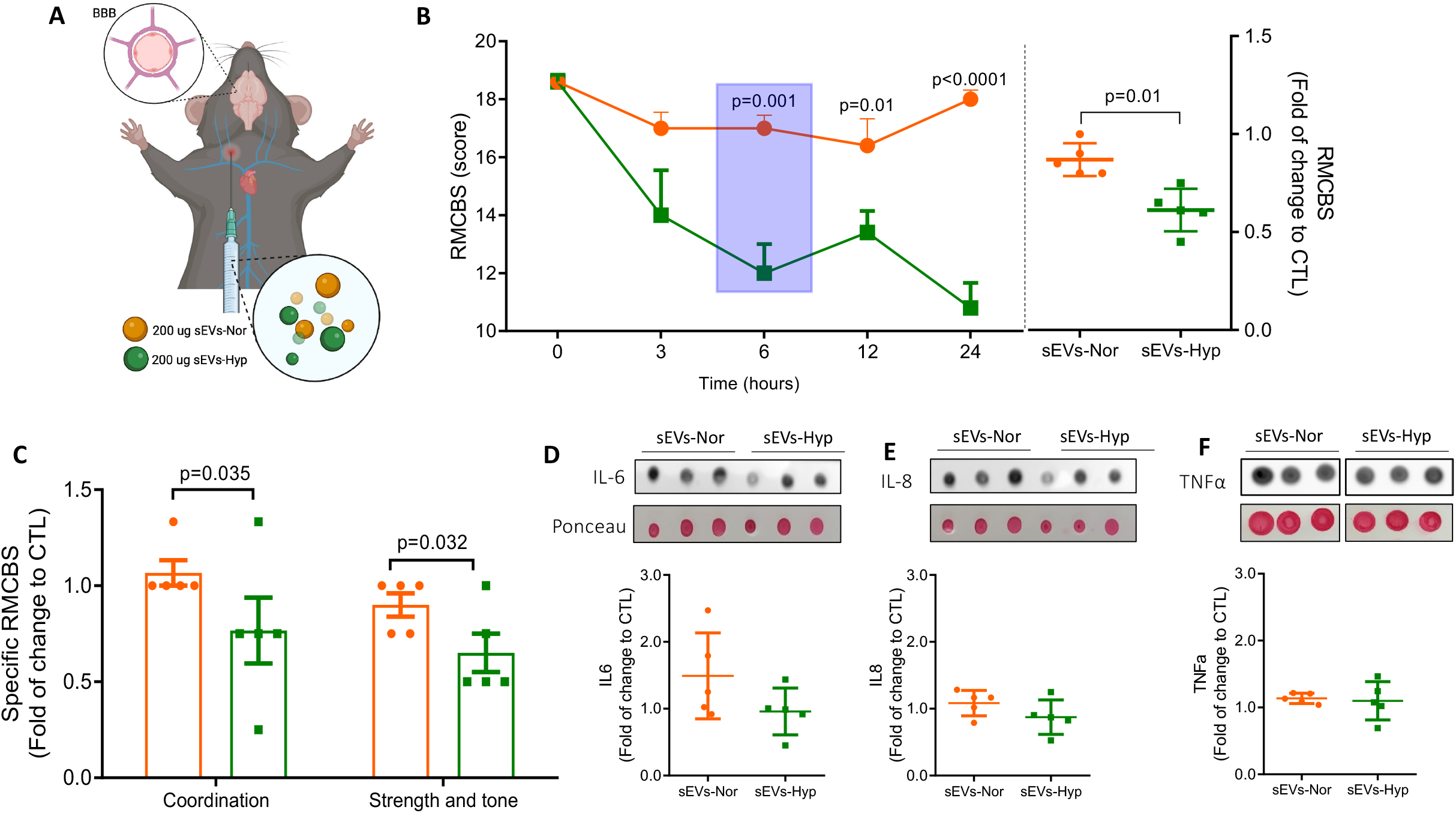
Effects of sEVs injection in non-pregnant mice. **A)** Cartoon of experimental in vivo approach in which sEVs (200 μg of total protein) from sEVs-Nor (orange dots) or sEVs-Hyp (green dots) extractions were injected into non-pregnant mice. **B)** RMCBS score in sEVs-Nor or sEVs-Hyp injected mice. The inserted chart indicated a fold change in the RMCBS score with respect to CTL. CLT mice have no changes in the RMCBS score (data not shown). **C)** Coordination and strength/tone parameters in the RMCBS score in mice injected as in B. **D)** IL-6, **E)** IL-8, and **F)** TNF-α levels expressed as fold of change with respect to CTL. Ponceau Red staining was used as a loading control. Every dot represents an individual sEV extraction from an individual patient. Data are presented as mean ± SD. P value is included when significant differences were found.

Non-pregnant mice injected with sEVs-Hyp showed an increased Evan’s blue extravasation compared with those injected with sEVs-Nor or control (i.e., injected with PBS) (Figure 4A). Evan’s blue extravasation in the former was 1.5-fold higher than sEVs-Nor (Figure 4B; P=0.007), particularly in the posterior sections (sections 7 to 9) of the analyzed brains (Figure 4C). sEVs-Nor and sEVs-Hyp (pre-incubated with a fluorescent dye, PKH67), were identified in brain blood vessels after 30 min of injection (Figure 4D). Disruption of the BBB generated with sEVs-Hyp administration was associated with a reduction in the protein levels of CLDN5 in most of the brain sections analyzed ex vivo (anterior-posterior; Figure 4E).

**Figure 4.**
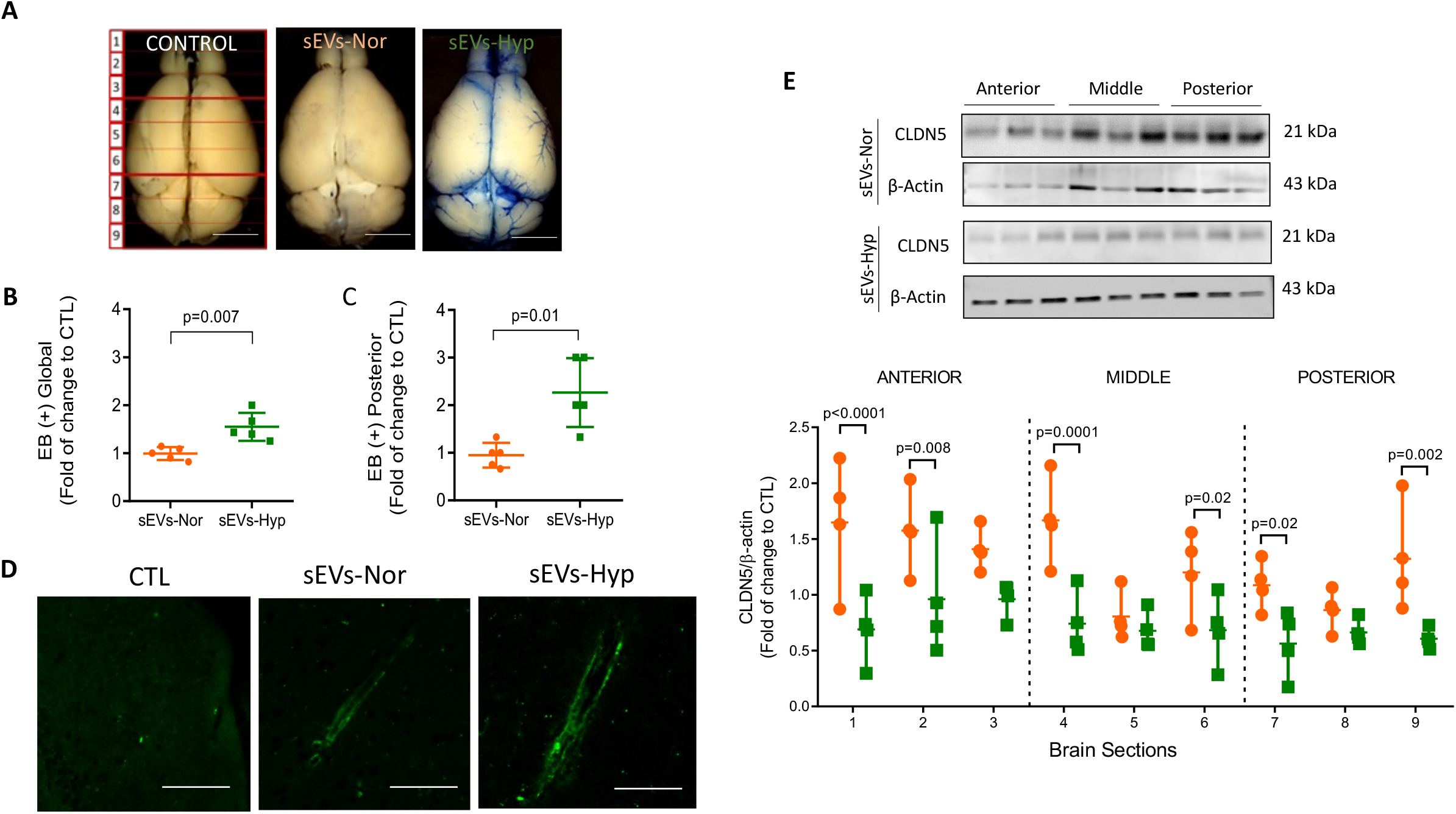
Evan’s blue extravasation and CLDN5 brain cortex levels in mice injected with sEVs. **A)** Representative image of control (CTL, injected with the vehicle); sEVs-Nor (orange dots) or sEVs-Hyp (green dots). After fixation, brains were cut into nine sections (1-9) anteroposterior. **B)** Estimation of Evan’s blue extravasation in the whole brain (global) or **C)** in the posterior area of the brain. **D)** Identification of sEVs-Nor and sEVs-Hyp in brain cortex (green fluorescence) 30 min after mice injection. CTL was injected with the vehicle. Picture magnification 100X. **E)** CLDN5 protein levels in each section of the brain cortex indicated in A. β-actin were used as loading control. The bottom charts indicate a densitometric analysis of the CLDN5/β-actin ratio. Values are expressed as a fold of change with respect to CTL. Every dot represents an individual sEV extraction from an individual patient. Data are presented as mean ± SD. P value is included when significant differences were found.

To explore the underlying molecular mechanisms behind sEVs-Hyp-induced disruption of the BBB, we estimated the content of VEGF, PlGF, and KDR due to their involvement in the disruption of the BBB^9,28-30^ (Figure 5A), or the VEGF-mediated reduction of CLND-5 in brain endothelial cells^30^. Remarkably, the sEVs content of VEGF was significantly higher in both sEVs-Hyp and sEVs-PE compared with sEVs-Nor or sEVs-NP, respectively (Figure 5D; P<0.05). Nevertheless, a high PlGF, but reduced KDR content, was found in sEVs-Hyp compared with sEVs-Nor. Yet no significant changes in these last proteins were found in sEVs-PE compared with sEVs-NP (Figure 5B, 5C, 5E, 5F).

**Figure 5.**
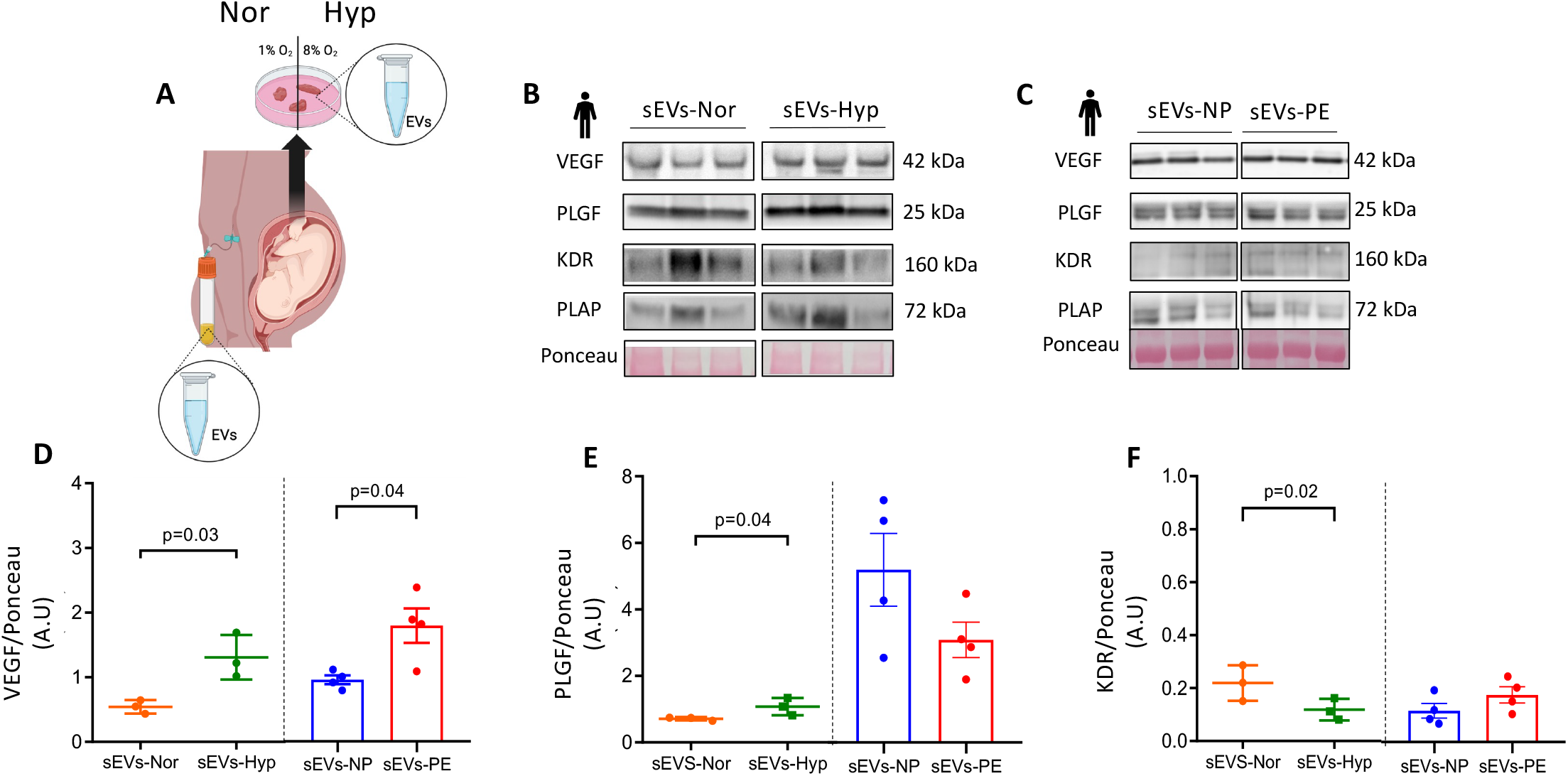
Analysis of VEGF-related proteins contends in sEVs. **A)** Cartoon showing experimental approach in which we collated sEVs-Nor (orange dots), sEVs-Hyp (green dots), and sEVs-NP (blue bars and dots) or sEVs-PE (red bars and dots) as indicated in Methods. **B)** Representative blots of VEGF, PlGF, KDR, and PLAP content in sEVs-Nor and sEVs-Hyp. **C)** or similar proteins investigated in sEVs-NP and sEVs-PE. Ponceau was used as a loading control. **D)** Densitometry analysis of VEGF/Ponceau ratio. **E)** PlGF/ Ponceau ratio. **F)** KDR/Ponceau ratio in the four experimental groups. Every dot represents an individual sEVs extraction from an individual patient. Data are presented as mean ± SD. P value is included when significant differences were found.

The potential involvement of activation (phosphorylation) of KDR receptors in the disruption of the BBB *in vitro* was analyzed. A significant reduction in the tyrosine 951 phosphorylation of KDR was found in cells exposed to sEVs-PE (Figure 6A-6C) or sEVs-Hyp (Figure 6D-6F), compared with those exposed to sEVs-NP or sEVs-Nor, respectively (Figure 6). Nevertheless, the total protein amount of KDR in hCMEC/D3 was significantly increased only in cells exposed to sEVs-PE compared with those treated with sEVs-NP.

**Figure 6.**
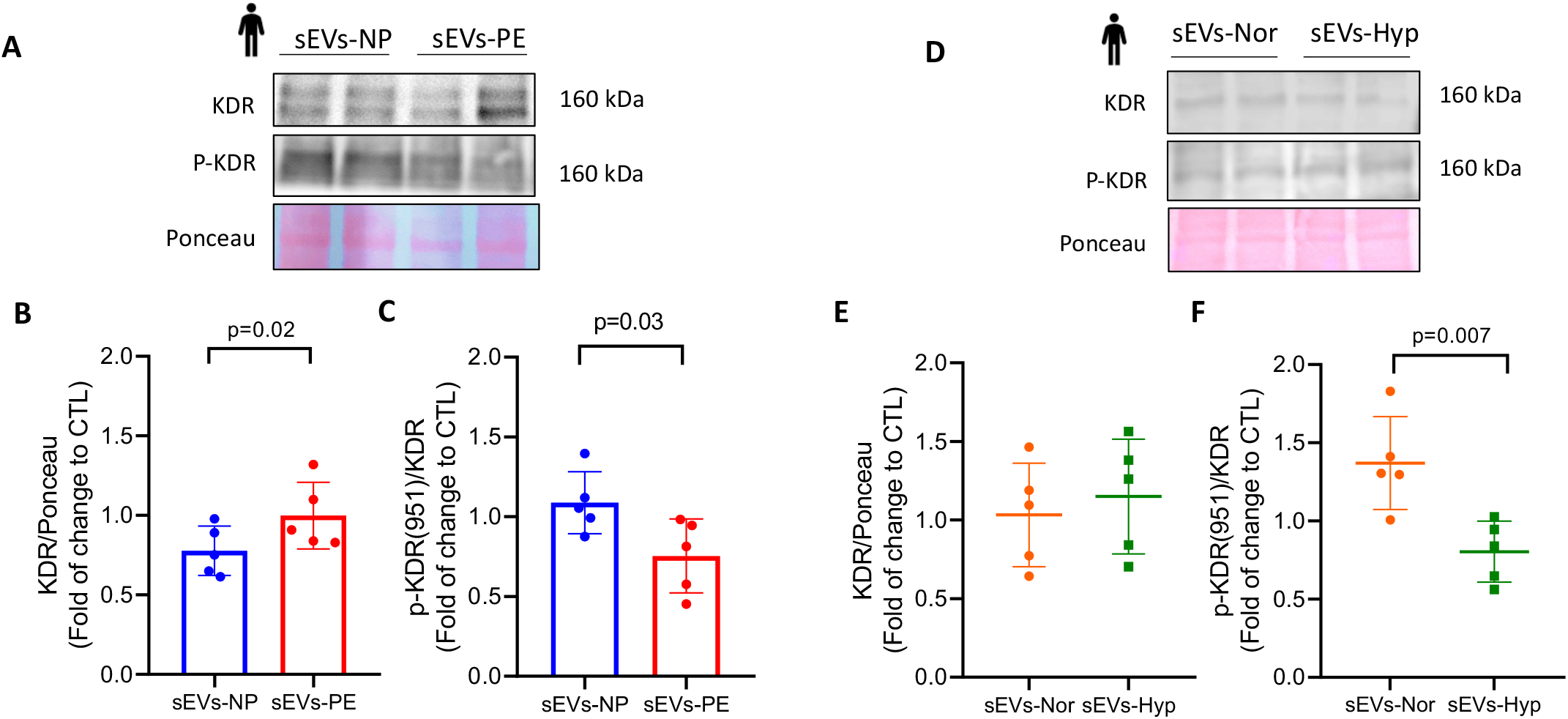
Analysis of KDR protein levels and activation (tyrosine 951 phosphorylation) in human brain endothelial cells treated with sEVs. **A and D)** Representative blot of KDR and KDR phosphorylated (P-KDR, tyrosine 951 phosphorylation) in human brain endothelial cells treated with sEVs-NP (blue bars and dots), sEVs-PE (red bars and dots), sEVs-Nor (orange dots) or sEVs-Hyp (green dots). **B and E)** Densitometry analysis of KDR/Ponceau or **C)** P-KDR/Ponceau ratio in human brain endothelium treated as in A. **E and F)** Densitometry analysis of KDR/Ponceau or P-KDR/Ponceau ratio in human brain endothelium is treated as in D. Every dot represents an individual sEVs extraction from an individual patient. Data are presented as mean ± SD. P value is included when significant differences were found.

## Discussion

In this study, we confirmed that sEVs-Hyp or sEVs-PE disrupt the BBB. Furthermore, sEVs-Hyp and sEVs-PE reduce the protein levels of CLDN5 in brain endothelial cells, suggesting that increased paracellular transport drives BBB disruption. Our *in vivo* studies indicate that sEVs-Hyp-mediated BBB disruption potentially constitutes a direct effect, since sEVs-Hyp was detected in the affected brain blood vessels and did not generate systemic inflammation, at least in the early time window (6 hours) when we found reduced mice’s cognition/functionality. Finally, our results suggest that VEGF content in sEVs-Hyp or sEVs-PE could act as the effector of CLDN5 reduction observed in brain endothelial cells. These results help us to understand the pathophysiology of cerebrovascular complications associated with PE.

Our findings constitute the first report showing a reduction in CLDN5 levels in cerebral endothelium as a potential underlying mechanism of BBB disruption generated by sEVs-PE (or sEVs-Hyp). It is widely known that CLDN5 is required for appropriate brain homeostasis, as observed in CLDN5^-/-^ mice pups, which die in the first 10 h postpartum, due to increased permeability and structural alterations in the conformation of the BBB^17^. In the human setting, CLDN5 is reduced in pathological conditions that exhibit disruption of the BBB, including stroke, traumatic brain edema, seizures, and psychiatric pathologies (i.e., anxiety or depression)^16^. In PE, our results demonstrated that maternal brains from the RUPP model show increased CLDN5 levels in the anterior area of the brain, which has no BBB disruption (i.e., Evan’s blue extravasation). In contrast, this up-regulation of CLDN5 was not observed in the posterior areas of the brain, which showed higher BBB disruption than their corresponding area in the brains of sham mice. We speculate that deregulation of CLDN5 could explain the occurrence of cerebral edema in the posterior area of the brain, as observed in our mice models (i.e., injected with sEVs-Hyp or RUPP). This finding is clinically relevant since the posterior area of the brain is more likely affected in women with eclampsia or posterior reversible encephalopathy syndrome (PRES)^2^.

Importantly, sEVs-PE or sEVs-Hyp also reduces the availability of CLDN5 in the cell membrane of brain endothelial cells. This effect occurs in the short term (3 and 6 h in both *in vitro* and *in vivo* models). Significantly, this reduction in CLDN5 depends on the integrity of the sEVs-PE. Multiple potential cellular mechanisms could explain this phenomenon, including, for example, that sEVs reduce the expression, cellular trafficking, or localization of CLDN5 on the cell membrane. This becomes more complex when we know that the half-life of CLDN5 is only 90 mins^31^. In this context, the content of microRNAs in sEVs-PE or sEVs-Hyp can block the expression of CLDN5. This possibility is feasible since our results indicate that sEVs-PE are taken up by endothelial cells, potentially delivering their cargo intracellularly. In addition, sonication impairs the uptake of sEVs-PE (data not shown), which is associated with preventing the reduction of CLDN5 in brain endothelial cells. Some reports have shown differential levels of miRNAs in sEVs-PE compared to sEVs-NP^32^. However, whether those miRNAs can downregulate the expression of CLDN5 needs further investigation.

miRNAs contained in sEVs can regulate the expression of target proteins in the recipient cells. In this regard, several miRNAs have been reported as blockers of the expression of CLDN5, including miR-21-3p, miR-30a, miR-101, miR-143, and miR-150^33^. Notably, high circulating levels of miR-21-3p in PE have been reported^34^, but no functional analysis of the potential transcript targets has been done so far. In addition to the miRNA’s regulation of target transcripts, it is also feasible that proteins contained in sEVs, such as VEGF (see below), upregulate the expression miRNAs in the target cell, which in turn can downregulate the CLDN5 mRNA levels. Therefore, we encourage future studies to investigate the miRNAs content in the sEVs-PE and sEVs-Hyp that targets CLDN5.

sEVs cargo also include functional proteins such as VEGF. Whether increased levels of VEGF observed in the sEVs-PE and sEVs-Hyp are directly involved in the disruption of the BBB is unknown. However, Argaw et al.,^30^ reported that VEGF downregulates the functional expression of CLDN5 in brain endothelial cells. VEGF-mediate downregulation of CLDN5 may involve the activation of the VEGF/VEGFR2/Pi3K-Akt pathway^35^. Accordingly, our results indicate that sEVs-PE increases protein levels of VEGF receptor 2 (VEGFR2 or KDR).

Our findings also indicate that sEVs-PE and sEVs-Hyp decrease the tyrosine 951 phosphorylation of KDR, suggesting a compensatory mechanism aimed at preventing/blocking the potential elevated activation of this receptor. Accordingly, we^9,36^ and others^29^ have reported that disruption of the BBB mediated by plasma of women with PE is prevented in the presence of inhibitors of KDR. We have reported that the plasma of women with PE increased the tyrosine 951 phosphorylation, which was associated with higher endothelial cell permeability^36^. sEVs may control KDR expression, but other plasma circulating factors may enhance KDR activation; we are testing this hypothesis in our laboratory.

Evidence included in this report enhances our understanding of the potential communication between the placenta and withing the brain vasculature via sEVs. In this regard, Han et al.,^37^, injected circulating placental-derived sEVs (PDsEVs) from women with PE into pregnant mice and found a significant reduction in cerebral blood flow shortly (seconds) after injection. Moreover, our group reported^7^ that administration of sEVs-Hyp, disrupts the BBB in non-pregnant mice. We extended these findings to show an identification of the sEVs-Nor and sEVs-Hyp in the brain vasculature after injection. Thus, we have confirmed that sEVs from the placenta can reach and modulate brain circulation. To further explore the potential effect of placental sEVs on brain function, we reported that sEVs-Hyp decreased the neurological evaluation (i.e., RMCBS) from 6-24 h after injection. Current areas of investigation in our group include understanding how the acute disruption of the BBB may have long-lasting consequences in the injected mice.

Placental sEVs are immunotolerant when injected into rodent *in vivo* models^37^. We extended these results to show that circulating proinflammatory molecules (IL-6, TNF-α, and IL-8) were unaffected after 6 h of placental sEV injection. However, they were increased at 24 h in the sEVs-Hyp injected mice. Despite that, our results suggest that disruption of the BBB may likely be associated with sEVs-Hyp direct effect over the BBB, since circulating proinflammatory cytokines were not affected despite a significant drop in the neurological evaluation observed in the injected mice.

Our study strengths include the combination of *in vivo, in vitro* (using human and mice brain endothelial cells), and *ex vivo* analysis of the disruption of the BBB generated with sEVs-Hyp and sEVs-PE. We also combine experiments with two animal models (non-pregnant, injected with sEVs and controls, and pregnant -sham and RUPP-mice), plus the inclusion of samples from women with and without PE. We combine the neurological evaluation of the animals with the analysis of the circulating and sEV cargo of critical proteins involved in the regulation of the BBB. However, we must also acknowledge some limitations. For instance, our findings regarding participation of VEGF/KDR needs further confirmation with inhibitors and with molecular systems in which we can study the sEVs delivery of these proteins into brain endothelial cells.

## Conclusion

sEV from hypoxic placentae disrupt the BBB, via direct interaction with brain endothelial cells associated with a reduction of CLDN5. This mechanism is correlated with a high concentration of VEGF in the placental sEVs-Hyp. The proposed mechanism can also be supported by sEVs-PE and by evidence in the PE-like syndrome in mice, suggesting that placental sEVs may damage the BBB in PE.

### Perspective

Cerebrovascular complications in women who experience preeclampsia have acute life-threatening consequences. Unfortunately, the underlying mechanisms of these complications are still unclear. However, preclinical, and clinical data indicate that these severe PE complications involve disruption of the BBB. This report confirms that sEVs isolated from plasma or placentae cultures in hypoxia can disrupt the BBB both *in vitro* and *in vivo*. Furthermore, we extend the knowledge by adding that disruption of the BBB may involve downregulating the critical tight junction CLDN5. We encourage future investigation to elucidate whether circulating placental sEVs or their cargo may be used as potential biomarkers for predicting cerebrovascular complications in women suffering from PE. Due to the relatively low incidence of cerebrovascular complications in high-income countries, this kind of analysis will need multicenter studies, including centers in LMIC. Since PE and its cerebrovascular complications are the leading cause of maternal death in LMIC, we believe that a better understanding of the pathophysiology of these complications demands the design of more focused ways to prevent and treat those deadly complications.

## Abbreviations

BBB: Blood-brain barrier
CBF: Cerebral blood flow
(CLDN5): claudin 5
FITC: Fluorescein-5-isothiocyanate
hCMEC/D3: human brain endothelial cell line,
BEND/3: mice brain endothelial cell line,
PE: Preeclampsia
PRES: Posterior reversible encephalopathy syndrome
sEVs: small extracellular vesicles or exosomes
sEVs-NP: sEVs from normal pregnancy
sEVs-PE: sEVs from preeclampsia
sEVs-Nor: sEVs from placentas cultures in normoxia
sEVs-Hyp: sEVs from placentas cultured in hypoxia
TEER: Transendothelial electrical resistance
TNF-α: Tumor necrosis factor-α
VEGF: Vascular endothelial growth factor
(KDR): VEGF receptor 2

## Acknowledgment

The authors would like to thank the researchers from GRIVAS Health.

## Funding

This study was funded by Fondecyt 1200250, and 1240295 (Chile) and HDM was funded under the terms of a BHF Basic Science Intermediate Basic Science Fellowship (FS/15/32/31604), UK Research and Innovation Grand Challenges Research Fund GROW Award scheme (grant number: MR/P027938/1) and NIHR–Wellcome Partnership for Global Health Research Collaborative Award (reference 217123/Z/19/Z).

## Author contributions

CE conceptualized the study and lead the research team. HS and BI performed most of the experiments in this manuscript. MC, FT, DC, LM, EEG, and JA perform selected experiments using both *in vitro* and *in vivo* approaches. HDM and LOK were responsible for patient recruitment. FOC and MV were consultants in exosome/sEVs characterization. CE, MV, and HDM edited the manuscript. All co-authors approved the final version of this manuscript.

## Disclosure

The authors do not have any conflict of interest to declare.

## Notes

**Conflict of interest:** none

### Competing Interest Statement

The authors have declared no competing interest.

